# Identification of genetic and non-genetic modifiers of genomic imprinting through whole genome screening in humans

**DOI:** 10.1101/2025.03.27.645693

**Authors:** Francesco Cecere, Raissa Relator, Michael Levy, Ankit Verma, Haley McConkey, Bruno Hay Mele, Laura Pignata, Carlo Giaccari, Emilia D’Angelo, Subham Saha, Abu Saadat, Angela Sparago, Claudia Angelini, Flavia Cerrato, Bekim Sadikovic, Andrea Riccio

## Abstract

Genomic imprinting is required for normal development, and abnormal methylation of differentially methylated regions (iDMRs) controlling the parent of origin-dependent expression of the imprinted genes has been found in constitutional disorders affecting growth, metabolism, neurobehavior, and in cancer. In most of these cases the cause of the imprinting abnormalities is unknown. Also, these studies have generally been performed on a limited number of CpGs, and a systematic investigation of iDMR methylation in the general population is lacking. By analysing the iDMRs in a large number of peripheral blood DNA methylation array datasets of unaffected individuals and patients with rare disorders, we determined the most common iDMR methylation profiles and identified many genetic and non-genetic factors contributing to their variability. We found that methylation variability is not homogeneous within the iDMRs and that the CpGs closer to the ZFP57 binding sites are less susceptible to methylation changes. We demonstrated the methylation polymorphism of three iDMRs and the atypical behaviour of several others, and reported the association of 25 disease- and 47 non-disease-complex traits, including blood cell type composition, as well as 15 mendelian or chromosomal disorders, with iDMR methylation changes in blood DNA. These findings identify several genetic and non-genetic factors associated with genomic imprinting maintenance in humans, which may have a role in the aetiology of the diseases associated with imprinting abnormalities and have clear implications in molecular diagnostics.

## Introduction

Genomic imprinting is a regulatory mechanism resulting in the mono-allelic and parent of origin- dependent expression of a few hundred mammalian genes (Barlow and Bartolomei 2014)). In its canonical form, imprinting results from differential establishment and maintenance of DNA methylation on the maternally and paternally derived autosomal chromosomes. More than fifty 1-2 kb-long regions that stably maintain differential DNA methylation (iDMRs) in multiple tissues, including blood, have been identified within the imprinted loci through the analysis of reciprocal genome-wide uniparental disomy (UPD) leukocyte samples (Court et al. 2014). The iDMRs include regions acquiring DNA methylation in the germline (gDMR) that are also defined imprinting control regions (ICRs) in that they control the expression of all the imprinted genes within a cluster through lncRNAs or other mechanisms. Other iDMRs, also known as somatic or secondary DMRs (sDMRs), are generally established on gene promoters during development, and their methylation depends on that of the closest gDMR.

Genomic imprinting is required for normal development and molecular defects affecting imprinted gene expression and function are associated with rare clinical disorders affecting growth, metabolism, endocrine and neuro-behavioral functions (Imprinting Disorders, ImpDis) (Eggermann et al. 2023). Typically, each ImpDis is caused by molecular abnormalities in a single locus, in many cases consisting of either hypo- or hyper-methylation of the locus-specific gDMR (Monk et al. 2019). In recent years, it has become evident that a subset of patients with ImpDis display mosaic multi-locus imprinting disturbance (MLID) that can be detected as loss of DNA methylation at one or more gDMRs in addition to that usually associated with the specific disease (Mackay et al. 2024). Although the clinical relevance of MLID is currently under investigation, this phenomenon appears to be of biological importance because several MLID cases display atypical clinical features of ImpDis, including high recurrence risk and maternal reproductive problems.

In some ImpDis cases, the methylation changes of the gDMRs are secondary to genetic variants (Monk et al. 2019). For example, either hypo- or hyper-methylation of a single gDMR can be caused by variants occurring in *cis,* while MLID can involve variants occurring in *trans*. The latter group includes *trans*-acting recessive variants of the transcription factor gene *ZFP57* in Transient Neonatal Diabetes and maternal-effect variants of the components of the oocyte Sub-Cortical Maternal Complex (SCMC) mainly in the Beckwith-Wiedemann syndrome (BWS) and Silver- Russell syndrome (Monk et al. 2019; Eggermann et al. 2022). Also, environmental factors may affect DNA methylation imprinting and an association between assisted reproduction technology (ART) and ImpDis has been reported by several studies (Monk et al. 2019). Nevertheless, the mechanisms underlying most iDMR methylation changes in ImpDis are unknown and their aetiology may be multifactorial (Monk et al. 2019).

The methylation abnormalities in ImpDis are often incomplete, suggesting that epigenetic mosaicism originated in early development (Monk et al. 2019). Diagnosis of ImpDis generally involves quantification of methylation levels at a few representative CpGs within a limited number of iDMRs relative to unaffected control baseline (Mackay et al. 2024). Thus, very little is known about the methylation levels of the iDMRs in the general population. The recent advancement in the use of genome-wide methylation arrays has allowed to accumulate large datasets of whole-genome DNA methylation profiles in control population and patients affected with rare disorders so that the methylation levels of thousands of CpGs can be studied in vast cohorts of individuals and eventually associated with complex or mendelian traits (Rooney and Sadikovic 2022; Levy et al. 2022; Xiong et al. 2022). Despite the availability of all this information, the methylation of the imprinted loci has not been systematically studied beyond the few ImpDis-related loci.

To investigate genome-wide maintenance of imprinted methylation, we analyzed the methylation array datasets obtained on blood DNA and studied the distribution of the iDMR methylation levels and their association with many factors in several thousand individuals, including patients affected by developmental disorders. The results describe the normal profile and variability of the iDMR CpG methylation and identify many genetic and non-genetic factors affecting it in humans.

## Results

### Methylation profiles of the iDMRs in the general population

We determined the methylation level of 49 human iDMRs (Supplemental Table S1) in the peripheral blood leukocyte DNA of a normal population by analyzing the methylation array datasets of 2664 individuals (Lehne et al. 2015; Wahl et al. 2017). Consistent with the differential methylation of their maternal and paternal alleles, most iDMRs displayed a mean methylation level close to 50% (Fig. 1a, Supplemental Table S2). Also, the distribution of their methylation levels generally resembled symmetric unimodal curves with a median value of 50% and was not strongly affected by the number of CpGs per iDMR assayed. However, certain loci deviated from this norm. For example, MKRN3:TSS, INPP5F:Int2 and GPR1-AS:TSS had median methylation levels well above 50%, and ZNF331:alt-TSS-DMR1 median methylation levels well below 50%. In addition, VTRNA2-1, WDR27:Int13 and IGF2r:Int2 showed multiple peaks, consistent with the polymorphic methylation described for one of these loci (Marttila et al. 2022; Romanelli et al. 2014). Other iDMRs, such as NNAT:TSS, DIRAS3:Ex2 and IGF1R:Int2, are more dispersed around the median indicating the presence of higher variability in the population.

**Fig. 1.**
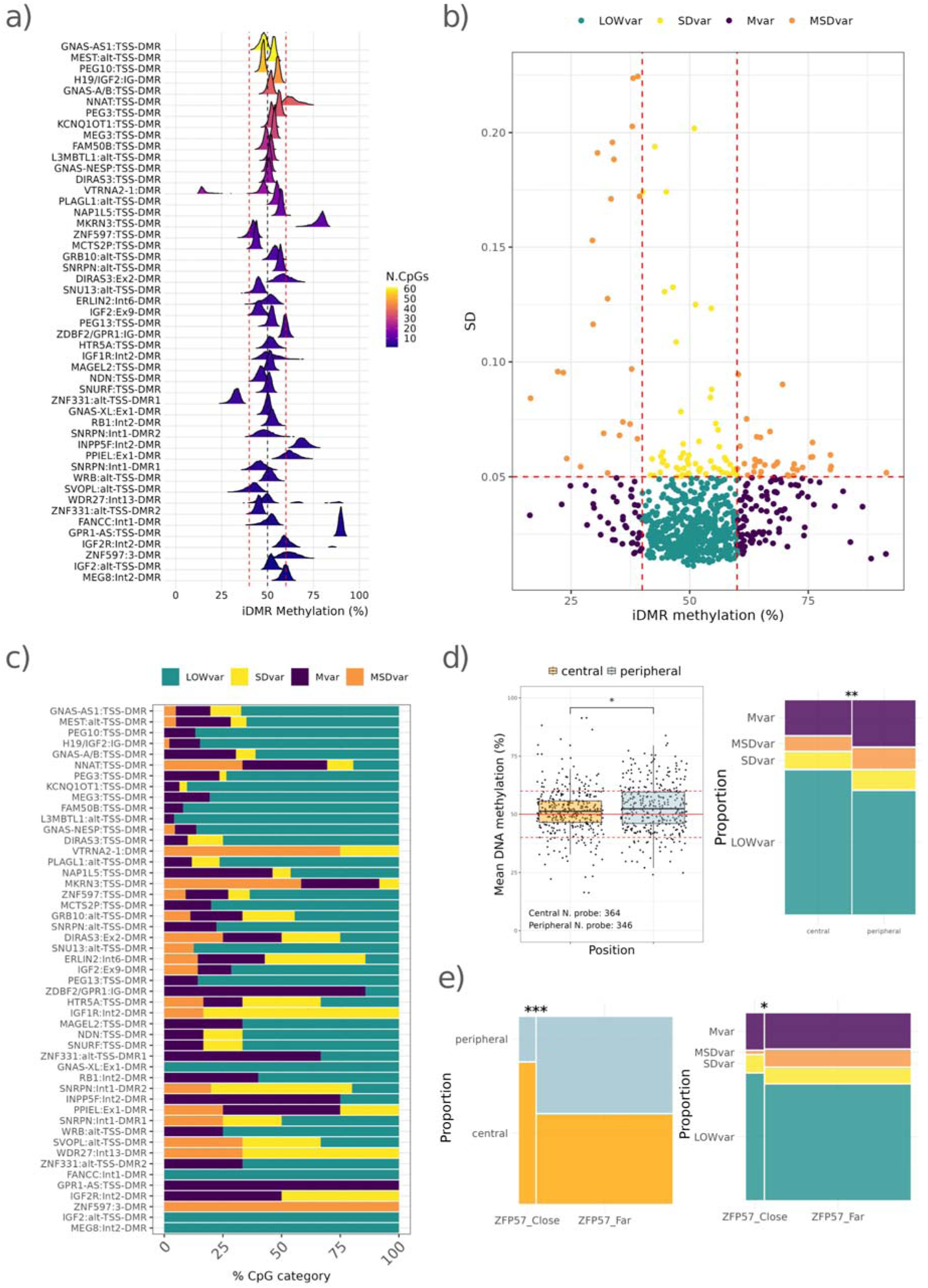
Methylation profiles of the iDMRs in the general population. **a)** Ridge plot showing the DNA methylation profile (βvalues) of 2,664 analyzed control samples across 49 iDMRs. The x-axis represents the DNA methylation value, while the y-axis lists the iDMRs. The density color indicates the number of probes covering each region. **b)** Scatterplot representing the relationship between the average DNA methylation value (x-axis) and the standard deviation (y-axis) for each iDMR CpG. Dots are color-coded based on variability thresholds defined by the red dashed lines: LOWvar (teal), Mvar (dark purple), SDvar (yellow) and MSDvar (orange). **c)** Stacked barplot showing the percentage distribution of LOWvar, Mvar, SDvar and MSDvar CpGs in each iDMR. **d)** iDMR subregion analysis: *Left*: Boxplot comparing the methylation values of the CpGs located in the peripheral versus central segments within the iDMRs. The significance is calculated using the Wilcoxson rank sum test. *Right*: Mosaic plot showing the proportion of SDvar, Mvar, MSDvar and LOWvar categories among the CpGs of the central or peripheral iDMR segments. The significance of the higher LOWvar-central probes is calculated using the Fisher-exact test. **e)** ZFP57 Peak Proximity: *Left*: Mosaic plot showing the proportion of the CpGs that are “Close” or “Far” from the ZFP57 peak summits in the peripheral and central iDMR subregions. The significance of the higher proportion of the CpGs “Close” to ZFP57 peaks in the central subregions is calculated using the Fisher-exact test. *Right*: Mosaic plot representing the proportion of SDvar, Mvar, MSDvar and LOWvar categories among the ZFP57 “Close” and “Far” CpGs. The significance of the higher proportion of the MSDvar among the CpGs “Far” from the ZFP57 peaks is calculated using the Fisher-exact test.

Our analysis covered 710 CpGs with 1-61 CpGs for each iDMR in the cohort under study (Fig. 1a, Supplemental Table S3). To determine how the individual CpGs influenced the distribution of iDMR methylation, we calculated the standard deviation (sd) and the mean methylation level of each iDMR CpG under the assumption of normal distribution of their methylation level. We identified 117 CpGs with sd>0.05 (SDvar CpGs) indicating high variability within the control population and 194 CpGs with mean methylation outside the [40%, 60%] range (Mvar CpGs) indicating atypical profiles. Of these, 58 CpGs met the criteria of both the SDvar and Mvar CpGs (MSDvar CpGs) (Fig.1b, Supplemental Table S4). We observed that VTRNA2-1, MKRN3:TSS, IGF1R:Int2, PPIEL:Ex1, WDR27:Int13, IGF2R:Int2 and ZNF597:3’ were enriched for both SDvar and Mvar CpGs, IGF1R:Int2 and SNRPN:Int1 were particularly enriched in SDvar CpGs, while ZDBF2/GPR1:IG, ZNF331:alt-TSS, INP55F:Int2 and GPR1-AS:TSS were particularly enriched in Mvar CpGs, (Fig. 1c and Supplemental Table S5). Apart from these, 457 CpGs (LOWvar CpGs) displayed sd<0.05 and mean methylation level between 40% and 60%, meeting the characteristics expected from imprinted loci. The LOWvar CpGs represented >50% of the CpGs in 29 iDMRs and overall were present in 41/49 loci (Fig. 1c). Indeed, when we reanalyzed the distribution of the iDMR mean methylation in our population after removing the SDvar CpGs, only the VTRNA2-1, WDR27:Int13, IGF1R:Int2 and ZNF597:3’ DMRs were completely lost, while the other iDMRs showed a more homogeneous profile (Supplemental Fig. S1a). On the other hand, when the Mvar CpGs were omitted, MEG8:Int2, GPR1-AS:TSS, INPP5F:Int2 and ZDBF2/GPR1 that strongly deviated from 50% methylation were lost, but the other iDMRs showed a mean methylation closer to 50% (Supplemental Fig. S1b). VTRNA1-1, WDR27:Int13 and IGF2R:Int2 represent an exception, in which the methylation level of all their CpGs are polymorphic. Finally, we obtained a methylation profile with a tiny peak close to 50% (Supplemental Fig. S1c) for 41/49 iDMRs, when only the LOWvar CpGs were considered.

To understand how CpG methylation was distributed within the iDMRs, we divided these regions into three segments: two peripheral and a central one. We observed that the methylation level of the central segments was closer to 50% with respect to the peripheral ones. Also, the peripheral segments had a higher number of Mvar and SDvar CpGs than the central ones, which instead were enriched in LOWvar CpGs (Fig. 1d, Supplemental Table S6), suggesting that the core of the iDMRs is more stable than their boundaries. This was particularly evident for some iDMRs (Supplemental Fig. S2a). To study the dependence of this trend on transcription factor binding, we intersected the coordinates of the central and peripheral iDMR segments with the binding sites (ChIPseq peaks) of ZFP57, which is a key protein in iDMR methylation maintenance (Quenneville et al. 2011; Imbeault et al. 2017). We found that the CpGs that are closer (<=250bp) to the summit of the ZFP57 peaks were more enriched in the central segments compared to the peripheral ones, while the CpGs that are more distant (>250bp) from the ZFP57 peaks were similarly distributed between peripheral and central segments (Fig. 1e, Supplemental Fig. 2b, Supplemental Table S7). Moreover, the CpGs that are more distant from the ZFP57 peaks were more represented by MSDvar than the CpGs that are closer to these binding sites (Fig. 1e, Supplemental Table S7).

In summary, two-thirds of the iDMR CpGs displayed a methylation level close to 50% with relatively low variability in blood DNA of a control population. However, a few iDMRs displayed highly variable CpG methylation and consistent deviation from 50% methylation level, including three polymorphic loci. Although variably methylated CpGs were present in the whole length of the iDMRs, these were less frequent in the central segments and in the vicinity of the ZFP57 binding sites.

### Association of blood iDMR methylation with complex traits

Once we assessed the variation of the iDMRs methylation level, we asked what factor affected the methylation of these regions within the general population. For this purpose, we retrieved the information deposited in the EWAS Data Hub and Atlas for the CpGs of the iDMRs. These data included high-quality associations of about 700 complex traits with CpG methylation in 3000 cohorts and 200 tissues/cells (Supplemental Table S8) (Xiong et al. 2022). For further analysis, we considered only the traits significantly associated with variation of iDMR methylation level in blood DNA. Also, we distinguished the traits into two categories: i. non-disease traits including environmental factors and phenotypic-behavioral traits; ii. disease traits, including neoplastic and non-neoplastic diseases. We found 107 iDMR CpGs associated with 47 non-disease traits and 111 iDMR CpGs associated with 25 disease traits (Fig. 2a, Supplemental Table S9). Overall, NNAT:TSS, VTRNA2, GNAS-AS1:TSS, MEST:alt-TSS and H19/IGF2:IG exhibited the highest number of CpGs associated with these traits (Fig 2b).

**Fig. 2.**
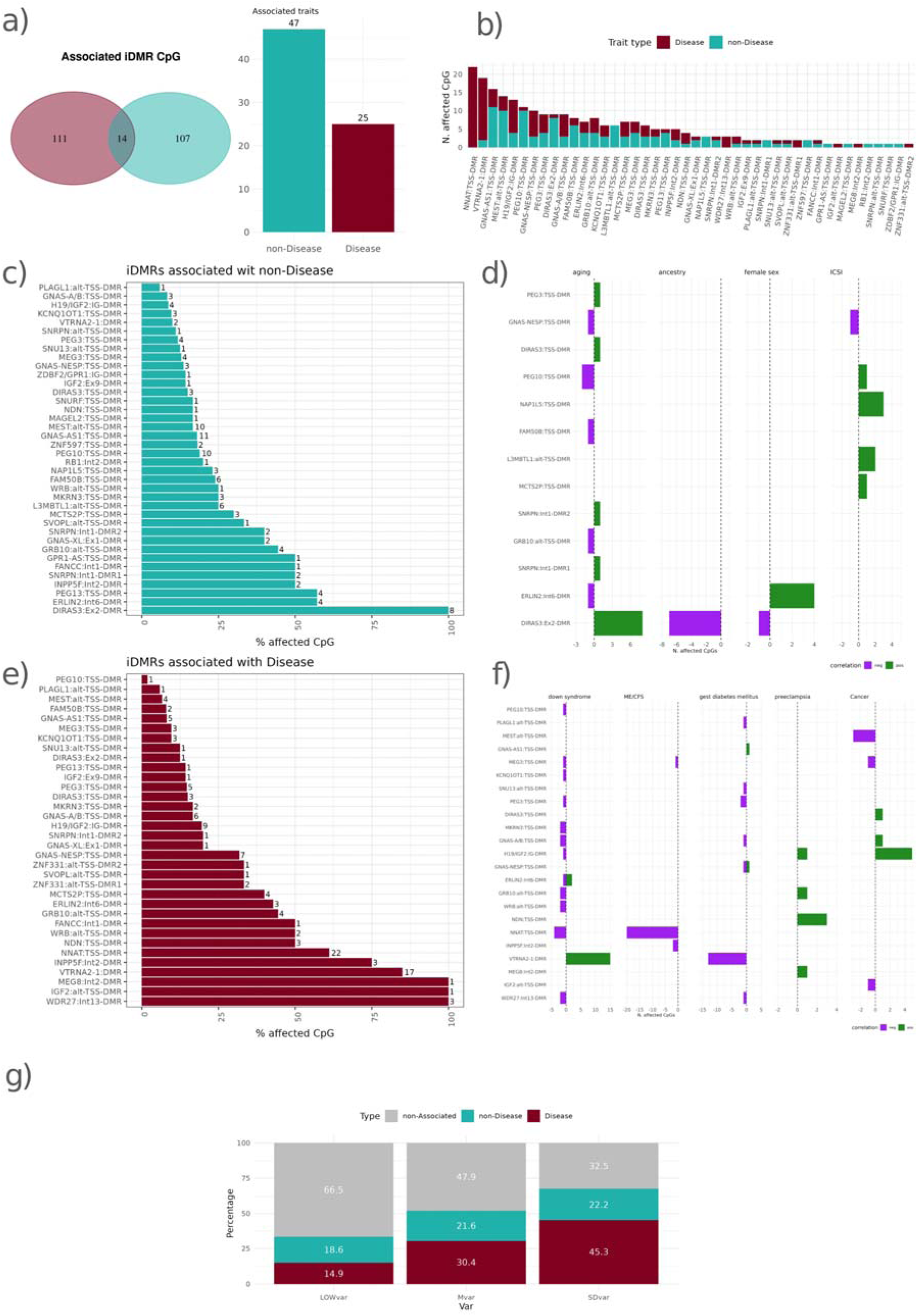
Association of complex traits with iDMR methylation. **a)** Associated traits with iDMR CpGs: *Left*: Venn diagram displaying the CpGs associated with Disease and non-Disease traits. *Right*: Barplot showing the number of unique disease (red) and non-disease (cyan) traits associated with iDMR methylation. **b)** Barplot showing the number of affected CpGs for each iDMR associated with disease (red) or non-disease (cyan) traits. **c)** Barplot representing the percentage of CpGs of each iDMR associated with non-disease traits. **d)** Barplot displaying the number of CpGs associated with specific non-disease traits. Negative correlations are in purple, positive correlations in forestgreen. **e)** Barplot representing the percentage of CpGs of each iDMR associated with Disease traits. **f)** Barplot displaying the number of CpGs associated with specific disease traits displayed as in *d*. **g)** Stacked Barplot showing the percentage of disease (red) and non-disease (cyan) trait associations with LOWvar, Mvar and SDvar CpGs.

DIRAS3:Ex2, ERLIN2:Int6 and PEG13:TSS were the iDMRs more strongly associated with the non-disease traits (Fig. 2c). In particular, the methylation of 8/8 DIRAS3:Ex2 CpGs were positively correlated with ageing and 7/8 of them also negatively correlated with European versus African ancestry (Fig. 2d). Ageing also affected fewer CpGs in several other iDMRs. In addition, 4/7 ERLIN2:Int6 CpGs were positively correlated with female sex (Fig. 2d). Furthermore, 30 CpGs distributed in 18 iDMRs (particularly FAM50B:TSS and MEST:alt-TSS) were associated with air pollution (Supplemental Fig. S3a). However, this effect was complex, depending on the nature of the pollutant examined (Additional file 2: Table S11). Furthermore, intracytoplasmic sperm injection (ICSI), a method of assisted reproduction technology, was associated with methylation changes of several iDMRs, particularly with 3 NAP1L5:TSS CpGs (Fig. 2d). Finally, maternal exposure to several potentially dangerous substances were associated with iDMR methylation. Particularly relevant is the negative correlation of maternal smoking with 3 MKRN3 CpGs and the negative correlation of several GNAS-AS1:TSS CpGs with prenatal perfluorooctanoate (PFOA) and perfluorohexane sulfonate (PFHxS) exposures (Supplemental Fig. S3a).

Apart from MEG8:Int2 and IGF2:alt-TSS with single CpGs covered, WDR27:Int13 (3/3), VTRNA2-1 (17/20), INPP5F:Int2 (3/4) and NNAT:TSS (22/36) were the iDMRs with the highest percentage of CpGs associated with disease-traits (Fig. 2e, Supplemental Table S11). In particular, the methylation of many iDMRs but VTRNA2-1, which was positively correlated, were negatively correlated with Down syndrome (Fig. 2f). Several iDMRs, including VTRNA2-1, were also correlated mostly negatively with gestational diabetes mellitus (Fig. 2f). Furthermore, we observed a negative correlation between myalgic encephalomyelitis/chronic fatigue syndrome (ME/CFS) and 20 NNAT:TSS CpGs (Fig. 2f), as well as a positive correlation between preeclampsia and 3 NDN:TSS CpGs (Fig. 2f). Blood DNA methylation of several DMRs were associated with cancer (Fig. 2f). In particular, we found 5 H19/IGF2:IG CpGs positively correlated with lung cancer, 3 MEST:alt-TSS CpGs negatively correlated with breast cancer, as well as the positive correlation of DIRAS3:TSS and GNAS-AB:TSS CpGs and the negative correlation of IGF2:alt-TSS and MEG3:TSS CpGs with further various cancer types (Supplemental Fig. S3b).

In order to investigate the relationship between the effect of the complex traits and the variability of iDMR methylation in the normal population, we intersected the information related to our control cohort and the association data (Supplemental Table S10). We found that more than 67% of the SDvar and 52% of the Mvar CpGs but only 33.5% of the LOWvar CpGs were associated with a trait (Fig. 2g). The prevalence of the Mvar and SDvar among the affected CpGs was more substantial for the disease-associated traits, but was evident for both trait categories. In summary, we found that iDMR methylation changes were associated with several environmental and complex traits and more concentrated on the SDvar and Mvar CpGs. These associations may explain part of the methylation variability observed in the general population.

### Blood cell type-specific methylation of the iDMRs

Blood is a tissue composed of many different cell types. A potential factor contributing to the variability of iDMR methylation levels in the human population could be ascribed to differences in blood cell-type composition (BCTC). To address this issue, we looked at the methylation level of the iDMR CpGs in the datasets of different blood cell types in the EWAS datahub (Supplemental Table S11). We found 63 iDMR CpGs whose methylation level had sd>0.05 across all the blood cell types (Fig. 3a). The BCTC effect was more substantial on NNAT:TSS and DIRAS3:Ex2 with 22 and 7 affected CpGs, respectively (Fig. 3b). In particular, the methylation level of NNAT:TSS was higher in T lymphocytes and primarily in CD127 regulatory and the CD45 memory T cells with respect to the other blood cell types, while that of DIRAS:Ex2 was higher in monocytes and neutrophils (Fig. 3c). Overall, 20/51 SDvar and 30/118 Mvar but only 25/432 LOWvar CpGs were affected by BCTC, with the highest number of SDvar and Mvar CpGs in NNAT:TSS and DIRAS3:Ex2 (Fig 3d). To investigate the effect of BCTC on their methylation profiles, we removed the affected CpGs from the NNAT:TSS and DIRAS3:Ex2 DMRs and plotted their methylation distribution in the control population. We observed that by using this filter the mean of the NNAT:TSS and DIRAS3:Ex2 distributions was closer to 50% (Fig. 3e). In summary, although BCTC affected the methylation level of a few CpGs in most iDMRs, NNAT:TSS and DIRAS3:Ex2 included a higher number of CpGs with different methylation in blood cell types and this may contribute to the high variability and deviation from the 50% level observed for these loci in the general population.

**Fig. 3.**
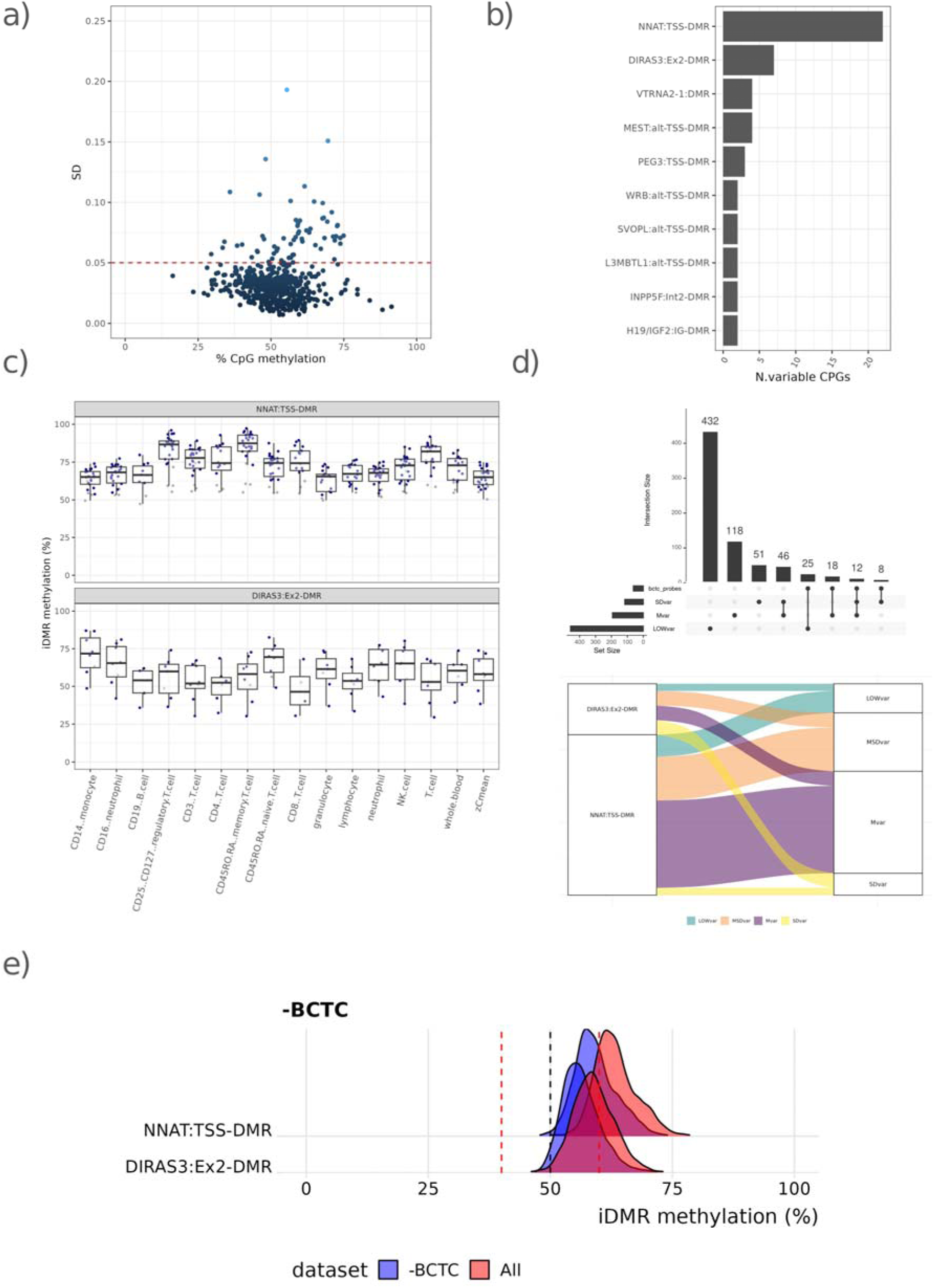
Effect of blood cell-type composition (BCTC) on iDMR methylation. **a)** Scatterplot cell types. The color scale of the dots indicates the SD value. **b)** Barplot displaying the number of iDMR CpGs affected by BCTC. **c)** Boxplot displaying the DNA methylation levels (βvalues) of the NNAT:TSS and DIRAS3:Ex2 DMRs in different blood cell types. The color scale of the dots (from dark grey to blue) indicates the strength of the BCTC effect (SD). **d)** Distribution of the BCTC- affected CpGs among the SDvar, Mvar, MSDvar and LOWvar categories. *Top*: Upset plot showing the intersection between SDvar, Mvar, MSDvar and LOWvar and the BCTC-affected CpGs. *Bottom*: Alluvial plot showing the proportions of SDvar, Mvar, MSDvar and LOWvar within the NNAT and DIRAS3:Ex2 iDMRs. **e)** Ridge plot showing the DNA methylation profiles (βvalues) of the DIRAS3:Ex2 and NNAT:TSS DMRs in the 2,664-individuals control cohort before and after the BCTC probes filtering. displaying the average values and standard deviation of iDMR methylation across different blood

### iDMR methylation in developmental disorders

Through the interrogation of a vast number of DNA methylation array datasets, methylation changes have been recently identified in the blood DNA of individuals affected by mendelian and chromosomal developmental disorders (DDs), and diagnostic episignatures have been identified in many cohorts (Rooney and Sadikovic 2022; Kerkhof et al. 2024). To identify genes that possibly affect genomic imprinting in humans, a screen for modifiers of iDMR methylation was conducted in samples of patients affected by 56 DDs that display mutations in the proteins of the epigenetic machinery and exhibit genome-wide DNA methylation episignatures within our EpiSign Knowledge Database (EKD) (Supplemental Table S12, see also https://episign.com/). Control samples (N=250) of healthy individuals and individuals negative for an existing EpiSign disorder were matched to DD cases using age, sex, and array type. It was observed that the matched controls exhibit the same trends seen in the general population, with most peaks near 50% methylation (Supplemental Fig. S4, Supplemental Table S13). Comparison of methylation status between EpiSign cohort cases and controls showed statistically significant (i.e., at least 10% mean difference and adjusted p-value < 0.05) difference in mean methylation levels for at least one iDMR in 15 cohorts (Table 1, Fig. 4a, and Supplemental Table S14).

**Fig. 4:**
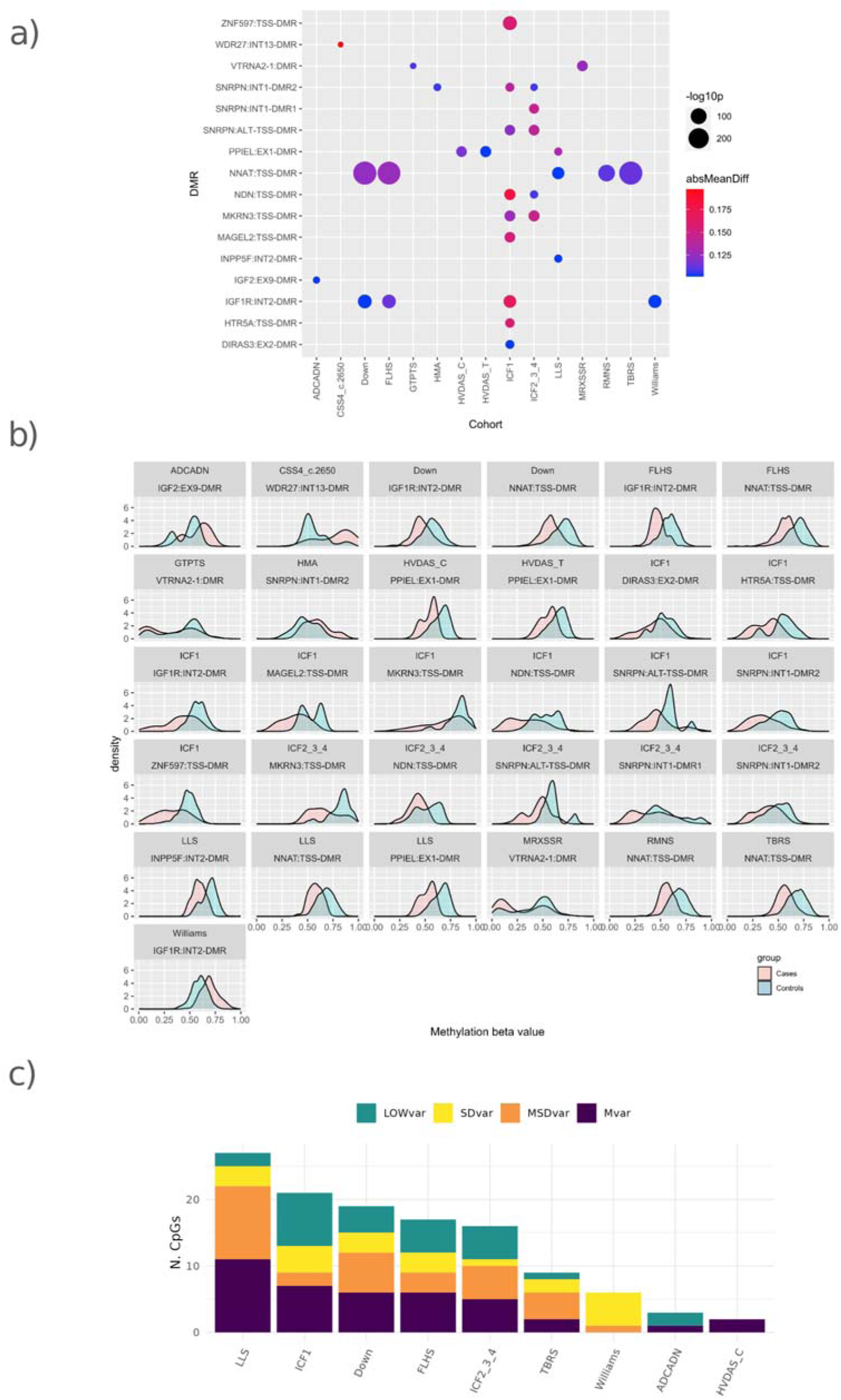
iDMR methylation changes in DDs. **a)** Bubble plot showing the differentially methylated iDMRs in the DD cohorts. The disorders are indicated on the x-axis and the iDMRs on the y-axis. The size of each bubble indicates the statistical significance (-log10 p-value), while the color represents the absolute mean methylation difference (absMeanDiff). **b)** Density plots comparing the methylation distributions of differentially methylated iDMRs between control individuals (blue) and DD patients (pink). **c)** Distribution of the CpGs with different methylation level in DDs vs controls among the SDvar, Mvar, MSDvar and LOWvar categories.

**Table 1.** Developmental disorders with affected iDMRs.

| Disorder | Gene(s) involved | Category | Patients number | Affected iDMRs |
| --- | --- | --- | --- | --- |
| Cerebellar ataxia, deafness, and narcolepsy, autosomal dominant (ADCADN) | *DNMT1* | DNA methyltransferase | 5 | IGF2:Ex9 |
| Coffin–Siris syndrome 4 (CSS4_c.2650) | *SMARCA4* | Chromatin remodeler | 3 | WDR27:Int13 |
| Down Syndrome (DS) | *HMGN1, MIS18A, AIRE, USP16, PRMT2, HLCS, DNMT3L, RBM11, CHAF1B, BRWD1* | Chromatin remodeler | 40 | IGF1R:Int2;<br>NNAT:TSS |
|  |  | Chromatin remodeler |  |  |
|  |  | Histone modification reader, Transcription factor |  |  |
|  |  | Histone ubiquitination |  |  |
|  |  | Histone methyltransferase |  |  |
|  |  | Histone modification writer |  |  |
|  |  | DNA methyltransferase |  |  |
|  |  | Alternative splicing |  |  |
|  |  | Chromatin remodeler |  |  |
|  |  | Histone modification reader |  |  |
| Floating-Harbour syndrome (FLHS) | *SRCAP* | Chromatin remodeler | 21 | IGF1R:Int2,<br>NNAT:TSS |
| Genitopatellar syndrome (GTPTS) | *KAT6B* | Histone acetyltransferase | 4 | VTRNA2-1:TSS |
| Helsmoortel–Van der Aa syndrome (ADNP syndrome (Central)) (HVDAS_C) | *ADNP* | Transcription factor | 14 | PPIEL:Ex1 |
| Helsmoortel–Van der Aa syndrome (ADNP syndrome (Terminal)) (HVDAS_T) | *ADNP* | Transcription factor | 21 | PPIEL:Ex1 |
| Hunter–McAlpine craniosynostosis syndrome (HMA) | *NPM1, UIMC1, NSD1* | Histone chaperone | 8 | SNRPN:Int1-DMR2 |
|  |  | Histone modification reader |  |  |
|  |  | Histone methyltransferase |  |  |
| Immunodeficiency, centromeric instability, facial anomalies syndrome 1 (ICF1) | *DNMT3B* | DNA methyltransferase | 8 | DIRAS3:Ex2,<br>HTR5A:TSS,<br>IGF1R:Int2, |
|  |  |  |  | MAGEL2:TSS,<br>MKRN3:TSS,<br>NDN:TSS,<br>SNRPN:alt-TSS,<br>SNRPN:Int1-DMR2,<br>ZNF597:TSS |
| Immunodeficiency, centromeric instability,<br>facial anomalies syndrome 2,3,4 (ICF2-3-4) | *ZBTB24*,<br>*CDCA7*, *HELLS* | Transcription factor and<br>chromatin remodelers | 7 | MKRN3:TSS,<br>NDN:TSS,<br>SNRPN:alt-TSS,<br>SNRPN:Int1-DMR1,<br>SNRPN:Int1-DMR2 |
| Intellectual developmental disorder, X-linked,<br>syndromic, Snyder–Robinson type<br>(MRXSSR) | *SMS* | Enzyme | 17 | VTRNA2-1:TSS |
| Luscan–Lumish syndrome (LLS) | *SETD2* | Histone<br>methyltransferase | 4 | INPP5F:Int2,<br>NNAT:TSS,<br>PPIEL:Ex1 |
| Rahman syndrome (RMNS) | *HIST1H1E* | Linker histone | 9 | NNAT:TSS |
| Tatton–Brown–Rahman syndrome (TBRS) | *DNMT3A* | DNA methyltransferase | 30 | NNAT:TSS |
| Williams–Beuren deletion syndrome<br>(Williams) | *BAZ1B*, *GTF2I* | Histone<br>phosphorylation<br>Transcription factor | 22 | IGF1R:Int2 |

In particular, 2-9 iDMRs were significantly deregulated in Immunodeficiency, centromeric instability, facial anomalies syndrome 1 (ICF1), Immunodeficiency, centromeric instability, facial anomalies syndrome 2,3,4 (ICF2,3,4), Luscan–Lumish syndrome (LLS), Down syndrome (DS) and Floating-Harbour syndrome (FLHS). Methylation of one iDMR was significantly altered in the remaining ten disorders, but for many of these, most of the other iDMRs followed a similar trend (Supplemental Table S14 and Supplemental Figs. S5-S60). Both gDMRs and sDMRs were affected, but some loci (e.g., NNAT, IGF1R, PPIEL and the SNRPN sDMRs) were more frequently deregulated than others (Additional file 2: Table S14). The mean iDMR methylation differences between DD patients and controls ranged from 10% to 20% (Fig. 4a and Additional file S2: Table S14), with an average of 13%, and the profiles of the patients and controls were partially overlapping (Fig. 4b). Furthermore, most of the cohort cases presented hypomethylated patterns compared to matched controls, except for the Cerebellar ataxia, deafness, and narcolepsy, autosomal dominant syndrome (ADCADN), Coffin–Siris syndrome 4 (CSS4_c.2650), Hunter– McAlpine craniosynostosis syndrome (HMA) and Williams-Beuren deletion syndrome (Williams) cohorts, which were hypermethylated in their respective deregulated DMRs (Fig. 4b).

We then looked at the methylation of the individual CpGs of the defective iDMRs in the DD cohorts and found 76 CpGs significantly deregulated in at least one disorder compared to the control cohort (Supplemental Table S15). According to their variability in the general population, we found 18 LOWvar, 14 Sdvar, 24 Mvar and 20 MSDvar CpGs deregulated in the DD patients (Fig. 4c). In summary, these results strongly suggest that the epigenetic regulators that are mutated in the DDs control the maintenance of iDMR methylation in blood DNA, with a stronger effect on the CpGs with moderate to high variability in the general population.

## Discussion

Defective iDMR methylation caused by unknown molecular mechanisms has been reported in ImpDis and cancer (Monk et al. 2019; Jelinic and Shaw 2007). However, the methodologies used to investigate the iDMRs in these studies were diverse and generally targeted a few CpGs. Although some evidence suggested polymorphic imprinting and interaction with environmental factors for some loci, the small sample size generally used in these studies limited information on iDMR methylation in the general population. On the other hand, high-throughput studies based on whole- genome methylation arrays have accumulated in the last few years and used to determine the association of environmental factors and complex and mendelian traits with DNA methylation changes. We employed this vast dataset to establish the iDMR methylation profiles in the general population and identified genetic and non-genetic factors contributing to the variability of these patterns in human blood DNA. We demonstrated that the methylation of about one-third of the iDMR CpGs deviates from the expected 50% level, is polymorphic or highly variable and identified several environmental and genetic factors contributing to this variability in the general population or in cohorts of individuals affected by developmental disorders.

Due to their gamete-of-origin-specific methylation, the iDMRs are expected to show a 50% methylation level in somatic cells. Although most of these regions displayed profiles compatible with these expectations, this was not the case for many loci when examined in a large population. In particular, the gDMRs ZNF331:alt-TSS1, INPP5F:Int2 and GPR1-AS:TSS and the sDMR MKRN3:TSS showed a mean methylation level different from 50%, indicating either *de novo* or loss of methylation in blood cells during development. Other iDMRs, such as VTRNA2-1:TSS, WDR27:Int13 and IGF2r:Int2 DMRs, displayed polymorphic methylation in our control cohort. These results are consistent with the reported polymorphic methylation imprinting of VTRNA2- 1:TSS and polymorphic expression imprinting of IGF2r:Int2 (Marttila et al. 2022; Romanelli et al. 2014; Monk et al. 2006; Xu et al. 1993). Other gDMRs, including NNAT:TSS, DIRAS3:Ex2, ERLIN2:Int6, PPIEL:Ex1, SVOPL:alt-TSS and ZNF597:3’, showed very high variability within our population. Also, methylation variability was not uniform within the iDMRs. The higher stability of the CpGs located in the vicinity of the ZFP57 binding sites that are more abundant in the center of the iDMRs indicate that transcription factor binding contributes to the maintenance of methylation imprinting in the general population.

Because blood cell types have diverse methylation level of many CpGs, changes in BCTC may influence the methylation profiles determined in blood DNA of an individual (Koestler et al. 2013). However, the effect of BCTC on iDMR methylation has not been investigated so far. We demonstrated that BCTC has little effect on most imprinted loci. However, NNAT:TSS and DIRAS:Ex2 are exceptions, in that the methylation level of the majority of their tested CpGs differ among blood cell types. This may contribute to their high sd and deviation from 50% methylation level observed in the general population and in some specific cohorts. For example, it is possible that the NNAT:TSS methylation changes we observed in DS are partly due to the frequent BCTC alterations these patients have (Zhang et al. 2023). Similarly, BCTC changes may underly at least in part the NNAT:TSS CpG hypomethylation detected in individuals affected by ME/CFS. Thus, it would be advisable to exclude these BCTC-affected CpGs when assaying methylation imprinting in diagnostic tests.

Genomic DNA methylation is deeply influenced by genetic and environmental factors (Xiong et al. 2022; Aref-Eshghi et al. 2020). The availability of many whole-genome array datasets, in which association with environmental conditions and complex traits was tested, prompted us to investigate to what extent iDMR methylation was affected by these relatively common factors. We found that many disease and non-disease-associated traits affect iDMR CpG methylation. Consistent with the dynamics of DNA methylation during human lifetime (Jones et al. 2015), ageing was the non-disease factor associated with more iDMRs. Although only one or two CpGs were significantly affected in most loci, this result demonstrates the importance of age-matched controls when measuring iDMR methylation in humans. Differently from the other iDMRs, many CpGs of DIRAS3:Ex2 were associated with ageing. These same CpGs were influenced by European versus African ancestry and BCTC. While the methylation differences between populations may be attributed to DNA sequence variants in the DIRAS3 locus (Husquin et al. 2018), the association with both ageing and BCTC suggests that methylation of this iDMR may be influenced by natural changes in the relative proportion of blood cell types in elder individuals (Marttila et al. 2025). Another interesting association is between ERLIN2:Int6 and the female sex. This DMR that is located within the imprinted Endoplasmic Reticulum Lipid Raft-Associated Protein 2 gene that is responsible for the *Spastic* Paraplegia 18A (Hedera 1993) is more methylated in girls compared to boys.

Several studies have reported an increased risk of developing ImpDis and particularly BWS in children conceived through Assisted Reproduction Technology, suggesting a disturbance in imprinting maintenance caused by the procedures used to enhance reproductive success (Ye et al. 2024). Consistent with these observations, we report an effect of intra-cytoplasmic sperm injection on several iDMRs. Also, air pollution had an effect on the methylation of several iDMRs. More specific associations were evident with prenatal exposure to smoking and the pollutants perfluorohexanesulfonic acid (PFHxS) and Perfluorooctanoic acid (PFOA). In particular, maternal smoking was strongly associated with hypomethylation of MKRN3:TSS, a locus involved in Central Precocious Puberty (Canton et al. 2025), while PFHxS and PFOA were associated with hypomethylation of GNAS-AS1:TSS, a molecular defect of Pseudohypoparathyroidism (Mantovani et al. 2018).

Trisomy 21 (Down syndrome, DS) has a widespread impact on DNA methylation, possibly caused by an imbalance of epigenetic regulator genes located in Chromosome 21 (Muskens et al. 2021). However, its consequence on genomic imprinting has not been investigated so far. We demonstrated hypomethylation of many iDMR CpGs in the blood DNA of DS patients. An elevated number of deregulated DMRs was also found in children exposed to gestational diabetes mellitus who are known to be at high risk of metabolic diseases (Weng et al. 2018). The affected PLAGL1, GNAS and PEG3 imprinted genes may have a role in the aetiology of these metabolic disturbances. The observed association of DS and gestational diabetes with VTRNA2-1 methylation should be taken with caution, because of the polymorphic imprinting of this locus. Indeed, VTRNA2-1 results can be explained by genetic association or bias in the frequency of allelic variants between patients and controls. Disturbed Placental Imprinting has been reported in preeclampsia (Wang et al. 2024). We found methylation changes of several iDMRs in blood leukocytes of women with early-onset preeclampsia, which may be explained by epigenetic modifications predisposing to this disease originated in early development. Finally, we demonstrated the association of iDMR methylation with several types of cancer. Although deregulated imprinting is a frequent finding in cancer cells, its alteration in blood leukocyte DNA has not been systematically investigated (Goovaerts et al. 2018; Cecere et al. 2023). The iDMR methylation changes we detected, particularly those of H19- IGF2:IG, IGF2:alt-TSS, MEST:alt-TSS and DIRAS3:TSS, may be originated in early development and represent markers for early tumour diagnosis.

Screening for DNA methylation modifiers by employing the array datasets of patients with developmental disorders harbouring mutations in proteins of the epigenetic machinery was successfully used a previous study of our group (Martin-Herranz et al. 2019). Using a similar approach, we identified several genes associated with iDMR methylation changes. First, we confirmed and expanded the results previously obtained in ICF1 and ICF2-4 (Velasco et al. 2018), which indicate the important role of DNMT3B and associated factors in imprinting maintenance. Among the new genes associated with iDMR methylation changes, we identified the one coding for the H3K36 methyltransferase SETD2 that in the mouse is required to establish imprinted methylation in oocytes (Xu et al. 2019). Further genes affecting iDMR methylation were those coding for the DNA methyltransferases DNMT1, the chromatin remodelers SRCAP and SMARCA4 and the transcription factor ADNP. Finally, the association of iDMR methylation changes with the HMA syndrome and the Williams syndrome that are characterized by the deletion of chromosome 17q23.1-q24.2 and 7q11.23, respectively, indicates that further epigenetic regulator genes controlling methylation imprinting maintenance are located in these regions. Concerning the associations between DDs and NNAT:TSS and VTRNA2-1 methylation, the considerations reported above for these two loci should be taken into accounts. Findings of disrupted DNA methylation profiles in iDMRs are consistent with and part of the broader, genome-wide disruption of DNA methylation profiles in rare neurodevelopmental disorders. Stability and reproducibility of these DNA methylation profiles across individuals affected by these disorders allow for development of highly accurate episignatures that can be used as diagnostic biomarkers in these disorders (Levy et al. 2022).

In summary, by looking at the methylation level of the imprinted loci in a large human cohort and the association of the imprinting methylation changes with many environmental and clinical variables, we have determined the CpGs of the imprinting control regions that are more resistant and those that change more frequently their methylation and demonstrated many genetic and non-genetic factors affecting their epigenetic variability. The faithful maintenance of genomic imprinting throughout cell division and differentiation is necessary for human health, but the genetic elements and factors that more stably maintain their epigenetic characteristics and the molecular mechanisms controlling this process are poorly defined. We hope this research will shed some light on these mechanisms and contribute to a better understanding of the aetiology of human diseases with disturbances of genomic imprinting.

## Methods

### Variability of iDMR methylation in human blood DNA

To determine the DNA methylation profiles of the iDMRs in the general population, we downloaded from GSE55763 the normalized Beta-values for each probe covered by the 450K Illumina methylation array across 2,711 samples. After identifying and removing technical replicates and cross-reacting probes (Chen et al. 2013), we retained only the probes covered across all samples, resulting in 398,094 probes and 2,664 samples for analysis. We obtained the iDMRs coordinates from the Human Imprinting Encyclopedia (http://www.humanimprints.net/) and used the bed_intersect function from the valr R package (v0.7.0) to filter the 710 probes that cover the iDMRs. We visualized the distribution of methylation levels across 49 iDMRs using ridge plots, while variability was assessed using standard deviation (SD) and mean methylation thresholds. Using this criteria, we classified the iDMR CpGs in LOWvar, Mvar, SDvar, and MSDvar CpGs based on their variability thresholds.

### Definition of the iDMR central and pheripheral segments

To determine the central and peripheral positions of the CpGs, we used a custom R script to divide the iDMRs genomic coordinates into three segments with a 1:2:1 proportion and with the central and peripheral parts covering 2/4 of the length each and including 364 and 346 probes, respectively. We used the information about the variability thresholds and performed a Fisher’s exact test to assess the enrichment of LOWvar, Mvar, SDvar, and MSDvar CpGs in central vs. peripheral regions.

### Evaluation of the effect of ZFP57 proximity on CpG methylation

We downloaded the ZFP57 ChIPseq peak summits from GSE200964 (de Tribolet-Hardy et al. 2023). Using a custom R script, we assigned the closest ZFP57 peak to each iDMR CpG. We defined “ZFP57_Close” CpGs those within 250 bp from the ChIPseq peak summit, and “ZFP57_Far” CpGs those located more than 250 bp from the ChIPseq peak summit. To evaluate the relationship between ZFP57 proximity and CpG variability, we performed a Fisher’s exact test to compare the distribution of MSDvar and LOWvar CpGs between ZFP57_Close and ZFP57_Far groups.

### EWAS analysis

To investigate the relationship between iDMR CpG methylation and complex traits, we retrieved trait association data from the EWAS Atlas (https://ngdc.cncb.ac.cn/ewas/atlas/index). We selected only the associations confirmed in blood DNA. Subsequently, intersecting the probeID of the CpGs covering the iDMRs, we retrieved the information on the traits associated with the methylation of the imprinted loci. To assess the impact of blood cell type composition (BCTC) on iDMR methylation, we downloaded the DNA methylation profiles of 25 blood cell types and identified 63 iDMR CpGs with a standard deviation (SD) > 0.05 across blood cell types.

### DNAm levels of DD patients and controls in imprinting regions

We compared levels of methylation of affected individuals with DDs and unaffected controls by employing our Episign Knowledge Database (EKD) DNA methylation data derived from peripheral blood (https://episign.com/). The EKD consists of cohorts for specific conditions as well as unaffected and healthy controls with varying age and gender, and type of array (Illumina 450k and EPIC). To ensure that the differences between methylation are due to genetic variation, we matched controls to cases by sex, age, and array type, if applicable. For each episignature cohort, we set the minimum number of controls to 150, and depending on the number of cases in a cohort, the case-control ratio ranges from 1:1 to 1:30. Beta values for probes in 50 iDMRs were investigated for both groups. Probes with missing values for at least 40% of samples in a group were removed. Missing values for the remaining probes are then imputed using the median value of that probe in the group. Ridge plots are generated to show density histogram of methylation beta values for each group in each signature cohort for every region. Furthermore, we calculated statistics to quantify methylation differences using two-sided z-test, with p-values adjusted to control for false discovery rate using the Bonferroni-Hochberg method, and population mean and standard deviation were computed using the data from matched controls. To identify differentially methylated probes (DMPs), we compared the methylation levels of syndromes to a control group using a z-score approach based on pooled variance .For each probe, we computed the pooled variance, which considers both the syndrome and control variances. Taking the square root gives the pooled standard deviation (SD). We then calculated z-scores to quantify how much the syndrome’s mean methylation differs from the control group, normalized by the pooled SD. To classify probes as differentially methylated, we set ±2 standard deviations (SDs) as threshold.

### Data access

Part of the DNA methylation data and metadata was obtained from the GEO public repository and are available under the following accession numbers: GSE55763 (control population) and GSE200964 (ZFP57 peaks summit). Individual genomic, epigenomic, or any other personally identifiable data that have not previously been made publicly available for samples in the EKD are prohibited from deposition in publicly accessible databases due to institutional and ethical restrictions.

### Ethics approval and consent to participate

All of the participants provided informed consent prior to sample collection. All of the samples and records were de-identified before any experimental or analytical procedures. The research was conducted in accordance with all relevant ethical regulations. All experimental methods comply with the Helsinki Declaration.

## Acknowledgements

This work was supported by grants from the Associazione Italiana Ricerca sul Cancro (AIRC; IG 2020 ID 24405), Fondazione Telethon (GMR23T1062), Italian Ministry of University and Research PRIN 2022B2N2BY awarded to AR, Progetto “National Centre for HPC, Big Data and Quantum Computing” finanziato nell’ambito del Piano Nazionale di Ripresa e Resilienza spoke 8 15858/2023 awarded to C.A., and Government of Canada through Genome Canada and the Ontario Genomics Institute (OGI-188) awarded to B.S. This project was conceptualized by B.S and A.R. Fr.C., R.R., B.S. and A.R. designed experiments and interpreted results. Fr.C., R.R., M.L., H.M.C., A.V., L.P., C.G., E.D’A., S.S., Ab.S., A.S., F.C., B.H.M. and C.A performed experiments. Fr.C., R.R., B.S. and A.R. wrote the manuscript with input from all authors. All authors reviewed the manuscript.

## References

1. Aref-Eshghi E, Kerkhof J, Pedro VP, Groupe DI France, Barat-Houari M, Ruiz-Pallares N, Andrau J-C, Lacombe D, Van-Gils J, Fergelot P, et al. 2020. Evaluation of DNA Methylation Episignatures for Diagnosis and Phenotype Correlations in 42 Mendelian Neurodevelopmental Disorders. Am J Hum Genet 106: 356–370.

2. Barlow DP, Bartolomei MS. 2014. Genomic imprinting in mammals. Cold Spring Harb Perspect Biol 6: a018382.

3. Canton APM, Macedo DB, Abreu AP, Latronico AC. 2025. Genetics and Epigenetics of Human Pubertal Timing: The Contribution of Genes Associated With Central Precocious Puberty. J Endocr Soc 9: bvae228.

4. Cecere F, Pignata L, Hay Mele B, Saadat A, D’Angelo E, Palumbo O, Palumbo P, Carella M, Scarano G, Rossi GB, et al. 2023. Co-Occurrence of Beckwith-Wiedemann Syndrome and Early-Onset Colorectal Cancer. Cancers 15: 1944.

5. Chen Y, Lemire M, Choufani S, Butcher DT, Grafodatskaya D, Zanke BW, Gallinger S, Hudson TJ, Weksberg R. 2013. Discovery of cross-reactive probes and polymorphic CpGs in the Illumina Infinium HumanMethylation450 microarray. Epigenetics 8: 203–209.

6. Court F, Tayama C, Romanelli V, Martin-Trujillo A, Iglesias-Platas I, Okamura K, Sugahara N, Simón C, Moore H, Harness JV, et al. 2014. Genome-wide parent-of-origin DNA methylation analysis reveals the intricacies of human imprinting and suggests a germline methylation-independent mechanism of establishment. Genome Res 24: 554–569.

7. de Tribolet-Hardy J, Thorball CW, Forey R, Planet E, Duc J, Coudray A, Khubieh B, Offner S, Pulver C, Fellay J, et al. 2023. Genetic features and genomic targets of human KRAB-zinc finger proteins. Genome Res 33: 1409–1423.

8. Eggermann T, Monk D, de Nanclares GP, Kagami M, Giabicani E, Riccio A, Tümer Z, Kalish JM, Tauber M, Duis J, et al. 2023. Imprinting disorders. Nat Rev Dis Primer 9: 33.

9. Eggermann T, Yapici E, Bliek J, Pereda A, Begemann M, Russo S, Tannorella P, Calzari L, de Nanclares GP, Lombardi P, et al. 2022. Trans-acting genetic variants causing multilocus imprinting disturbance (MLID): common mechanisms and consequences. Clin Epigenetics 14. https://clinicalepigeneticsjournal.biomedcentral.com/articles/10.1186/s13148-022-01259-x (Accessed March 17, 2025).

10. Goovaerts T, Steyaert S, Vandenbussche CA, Galle J, Thas O, Van Criekinge W, De Meyer T. 2018. A comprehensive overview of genomic imprinting in breast and its deregulation in cancer. Nat Commun 9: 4120.

11. Hedera P. 1993. Hereditary Spastic Paraplegia Overview. In *GeneReviews®* (eds. M.P. Adam, J. Feldman, G.M. Mirzaa, R.A. Pagon, S.E. Wallace, and A. Amemiya), University of Washington, Seattle, Seattle (WA) http://www.ncbi.nlm.nih.gov/books/NBK1509/ (Accessed March 17, 2025).

12. Husquin LT, Rotival M, Fagny M, Quach H, Zidane N, McEwen LM, MacIsaac JL, Kobor MS, Aschard H, Patin E, et al. 2018. Exploring the genetic basis of human population differences in DNA methylation and their causal impact on immune gene regulation. Genome Biol 19: 222.

13. Imbeault M, Helleboid P-Y, Trono D. 2017. KRAB zinc-finger proteins contribute to the evolution of gene regulatory networks. Nature 543: 550–554.

14. Jelinic P, Shaw P. 2007. Loss of imprinting and cancer. J Pathol 211: 261–268.

15. Jones MJ, Goodman SJ, Kobor MS. 2015. DNA methylation and healthy human aging. Aging Cell 14: 924–932.

16. Kerkhof J, Rastin C, Levy MA, Relator R, McConkey H, Demain L, Dominguez-Garrido E, Kaat LD, Houge SD, DuPont BR, et al. 2024. Diagnostic utility and reporting recommendations for clinical DNA methylation episignature testing in genetically undiagnosed rare diseases. Genet Med Off J Am Coll Med Genet 26: 101075.

17. Koestler DC, Christensen BC, Karagas MR, Marsit CJ, Langevin SM, Kelsey KT, Wiencke JK, Houseman EA. 2013. Blood-based profiles of DNA methylation predict the underlying distribution of cell types. Epigenetics 8: 816–826.

18. Lehne B, Drong AW, Loh M, Zhang W, Scott WR, Tan S-T, Afzal U, Scott J, Jarvelin M-R, Elliott P, et al. 2015. A coherent approach for analysis of the Illumina HumanMethylation450 BeadChip improves data quality and performance in epigenome-wide association studies. Genome Biol 16: 37.

19. Levy MA, McConkey H, Kerkhof J, Barat-Houari M, Bargiacchi S, Biamino E, Bralo MP, Cappuccio G, Ciolfi A, Clarke A, et al. 2022. Novel diagnostic DNA methylation episignatures expand and refine the epigenetic landscapes of Mendelian disorders. Hum Genet Genomics Adv 3: 100075.

20. Mackay DJG, Gazdagh G, Monk D, Brioude F, Giabicani E, Krzyzewska IM, Kalish JM, Maas SM, Kagami M, Beygo J, et al. 2024. Multi-locus imprinting disturbance (MLID): interim joint statement for clinical and molecular diagnosis. Clin Epigenetics 16. https://clinicalepigeneticsjournal.biomedcentral.com/articles/10.1186/s13148-024-01713-y (Accessed March 17, 2025).

21. Mantovani G, Bastepe M, Monk D, de Sanctis L, Thiele S, Usardi A, Ahmed SF, Bufo R, Choplin T, De Filippo G, et al. 2018. Diagnosis and management of pseudohypoparathyroidism and related disorders: first international Consensus Statement. Nat Rev Endocrinol 14: 476–500.

22. Martin-Herranz DE, Aref-Eshghi E, Bonder MJ, Stubbs TM, Choufani S, Weksberg R, Stegle O, Sadikovic B, Reik W, Thornton JM. 2019. Screening for genes that accelerate the epigenetic aging clock in humans reveals a role for the H3K36 methyltransferase NSD1. Genome Biol 20: 146.

23. Marttila S, Rajić S, Ciantar J, Mak JKL, Junttila IS, Kummola L, Hägg S, Raitoharju E, Kananen L. 2025. Biological aging of different blood cell types. GeroScience 47: 1075–1092.

24. Marttila S, Tamminen H, Rajic S, Mishra PP, Lehtimaki T, Raitakari O, Kähönen M, Kananen L, Jylhävä J, Hägg S, et al. 2022. Methylation status of VTRNA2-1/nc886 is stable across populations, monozygotic twin pairs and in majority of tissues. Epigenomics 14: 1105–1124.

25. Monk D, Arnaud P, Apostolidou S, Hills FA, Kelsey G, Stanier P, Feil R, Moore GE. 2006. Limited evolutionary conservation of imprinting in the human placenta. Proc Natl Acad Sci U S A 103: 6623–6628.

26. Monk D, Mackay DJG, Eggermann T, Maher ER, Riccio A. 2019. Genomic imprinting disorders: lessons on how genome, epigenome and environment interact. Nat Rev Genet 20: 235–248.

27. Muskens IS, Li S, Jackson T, Elliot N, Hansen HM, Myint SS, Pandey P, Schraw JM, Roy R, Anguiano J, et al. 2021. The genome-wide impact of trisomy 21 on DNA methylation and its implications for hematopoiesis. Nat Commun 12: 821.

28. Quenneville S, Verde G, Corsinotti A, Kapopoulou A, Jakobsson J, Offner S, Baglivo I, Pedone PV, Grimaldi G, Riccio A, et al. 2011. In embryonic stem cells, ZFP57/KAP1 recognize a methylated hexanucleotide to affect chromatin and DNA methylation of imprinting control regions. Mol Cell 44: 361–372.

29. Romanelli V, Nakabayashi K, Vizoso M, Moran S, Iglesias-Platas I, Sugahara N, Sugahara N, Simón C, Simón C, Hata K, et al. 2014. Variable maternal methylation overlapping the nc886/vtRNA2-1 locus is locked between hypermethylated repeats and is frequently altered in cancer. Epigenetics 9: 783–790.

30. Rooney K, Sadikovic B. 2022. DNA Methylation Episignatures in Neurodevelopmental Disorders Associated with Large Structural Copy Number Variants: Clinical Implications. Int J Mol Sci 23: 7862.

31. Velasco G, Grillo G, Touleimat N, Ferry L, Ivkovic I, Ribierre F, Deleuze J-F, Chantalat S, Picard C, Francastel C. 2018. Comparative methylome analysis of ICF patients identifies heterochromatin loci that require ZBTB24, CDCA7 and HELLS for their methylated state. Hum Mol Genet 27: 2409–2424.

32. Wahl S, Drong A, Lehne B, Loh M, Scott WR, Kunze S, Tsai P-C, Ried JS, Zhang W, Yang Y, et al. 2017. Epigenome-wide association study of body mass index, and the adverse outcomes of adiposity. Nature 541: 81–86.

33. Wang X, Liu Y, Wu Y, Lin C, Yang S, Yang Y, Chen D, Yu B. 2024. Methylation alterations of imprinted genes in different placental diseases. Clin Epigenetics 16: 132.

34. Weng X, Liu F, Zhang H, Kan M, Wang T, Dong M, Liu Y. 2018. Genome-wide DNA methylation profiling in infants born to gestational diabetes mellitus. Diabetes Res Clin Pract 142: 10–18.

35. Xiong Z, Yang F, Li M, Ma Y, Zhao W, Wang G, Li Z, Zheng X, Zou D, Zong W, et al. 2022. EWAS Open Platform: integrated data, knowledge and toolkit for epigenome-wide association study. Nucleic Acids Res 50: D1004–D1009.

36. Xu Q, Xiang Y, Wang Q, Wang L, Brind’Amour J, Bogutz AB, Zhang Y, Zhang B, Yu G, Xia W, et al. 2019. SETD2 regulates the maternal epigenome, genomic imprinting and embryonic development. Nat Genet 51: 844–856.

37. Xu Y, Goodyer CG, Deal C, Polychronakos C. 1993. Functional polymorphism in the parental imprinting of the human IGF2R gene. Biochem Biophys Res Commun 197: 747–754.

38. Ye M, Reyes Palomares A, Iwarsson E, Oberg AS, Rodriguez-Wallberg KA. 2024. Imprinting disorders in children conceived with assisted reproductive technology in Sweden. Fertil Steril 122: 706–714.

39. Zhang Z, Stolrow HG, Christensen BC, Salas LA. 2023. Down Syndrome Altered Cell Composition in Blood, Brain, and Buccal Swab Samples Profiled by DNA-Methylation-Based Cell-Type Deconvolution. Cells 12: 1168.

